# Real Bodies Not Required? Placebo Analgesia and Pain Perception in Immersive Virtual and Augmented Reality

**DOI:** 10.1101/2020.12.18.423276

**Authors:** Jasmine T. Ho, Peter Krummenacher, Marte Roel Lesur, Bigna Lenggenhager

**Affiliations:** University of Zurich, Department of Psychology, Binzmühlestrasse 14, Box 9, 8050 Zurich, Switzerland; Brainability, LLC, Sonneggstrasse 86, 8006 Zurich, Switzerland; Experimental and Clinical Pharmacopsychology, Department of Psychiatry, Psychotherapy, Psychotherapy and Psychosomatics, 8032 Zurich, Switzerland; University of Zurich, Department of Psychology

**Author notes:** All authors have seen and approved the manuscript, which has not been accepted or published elsewhere. No competing interests exist.

**Keywords:** Placebo analgesia, virtual placebo, virtual reality, augmented reality, virtual embodiment, expectation, pain

## Abstract

Pain represents an embodied process, wherein inferences are not only drawn from sensory inputs, but also from bodily states. Previous research has demonstrated that a placebo administered to an embodied rubber hand can effectively induce analgesia, providing first evidence that placebos can work even when applied to temporarily embodied, artificial body parts. Using a heat pain paradigm, the present study investigates placebo analgesia and pain perception during virtual embodiment. We examined whether a virtual placebo (a sham heat protective glove) can successfully induce analgesia, even when administered to a virtual body. The analgesic efficacy of the virtual placebo to the real hand (augmented reality setting) or virtual hand (virtual reality setting) was compared to a physical placebo administered to the own, physical body (physical reality setting). Furthermore, pain perception and subjective embodiment were compared between settings. Healthy participants (n=48) were assigned to either an analgesia-expectation or control-expectation group, where subjective and objective pain was measured at pre- and post-intervention time points. Results evinced that pre-intervention pain intensity was lower in the virtual reality setting, and that participants in the analgesia-expectation condition, after the intervention, exhibited significantly higher pain thresholds, and lower pain intensity and unpleasantness ratings than control-expectation participants, independent of the setting. Our findings evince that a virtual placebo can elicit placebo analgesia comparable to that of a physical placebo, and that administration of a placebo does not necessitate physical bodily interaction to produce analgesic responses, which might pave the way for effective new non-pharmacological approach for pain management.

## 1. Introduction

Placebo processes, during which inert substances or sham treatments produce behavioral and physiological changes such as symptom improvement, represent a topic of stimulating scientific inquiry [2,4,54]. Placebo analgesia constitutes one of the most studied and best understood placebo responses, and has been shown to be mediated by expectations, which in turn can be moderated by forms of learning such as classical conditioning, verbal cues, environmental cues (e.g., white lab coat or hospital settings) or social observation [11,12].

Many processes that have been the focus of placebo research, such as pain [5,13,56,66], can be considered embodied processes, wherein inferences are not only drawn from sensory inputs, but also from bodily states [18,43]. The experience of pain represents a context-dependent, multidimensional process that depends on the interplay between sensory and cognitive-affective mechanisms, and has shown to be influenced by the subjective sense of embodiment [35,52]. Experimentally induced alterations of embodiment have demonstrated efficacy in modulating both acute [23,34,35] and chronic [49,60] pain. For example, it has been shown that color modification of an embodied virtual arm through a red color sensitizes participants to thermal stimulations [36], while increasing the size of a viewed hand was able to reduce pain evinced by increased heat-pain thresholds [32]. An out-of-body illusion successfully reduced pain in various different types of chronic pain conditions [47], while body illusions aimed at reducing body perception disturbances in osteoarthrits (i.e., stretching of the painful limb) also led to significant reductions in pain [60], some of which experienced extended analgesia lasting hours or even weeks after only one session [48]. Morevover, a recent rubber hand illusion (RHI) paradigm, where synchronous stroking of the own (hidden from view) arm and a viewed rubber hand creates an illusory ownership of the rubber hand, demonstrated that not only basic pain perception [16] but also placebo analgesia [9] depends on the sense of embodiment. Such body illusions constitute representative experimental paradigms to explore the potentially moderating role of embodiment (i.e., how we experience the own body) in placebo processes.

Immersive virtual reality (VR) and augmented reality (AR) technology has facilitated illusory alterations of embodiment, where entire physical bodies can be substituted with virtual surrogates [58]. The strong sense of presence [53] elicited by immersive VR enables realistic simulations, where multisensory interactions with the virtual body can elicit autonomic responses and motor cortex activations equivalent to real-world experiences [57]. Therefore, emobdied virtual reality might offer a promising area for future translational non-pharmacological pain management tools. However, a pertinent difference between these VR and AR modalities concerns the level of embodiment experienced in each: Whereas AR can retain physical components, such as one’s own body, VR introduces complete virtual surrogates that visually replace the own physical body. By employing multisensory stimulation, such as synchronous visuomotor feedback between the virtual avatar’s and one’s own body’s movements induces a sense of embodiment of the virtual body [37]. As a foundational basis of our self, both the holistic sense of (bodily) self, as well as the individual components that make up our sense of embodiment (i.e., the sense of agency, location, ownership, and perspective) constitute a continuous source of information that we use for the interaction with our environmental surroundings. To investigate the influence of physical versus virtual embodiment, we employed embodied VR and AR to study the influence of embodiment in different settings (VR, AR, physical reality (PR)) on placebo responses and pain perception. The present study bridges the domains of placebo analgesia and virtual embodiment by examining whether a virtual placebo (in VR and AR) can successfully induce analgesia, even when it is administered to only a virtual body (VR). Furthermore, we compared the analgesic efficacy of these open label (i.e., placebos that entail no deception surrounding their inert nature) virtual placebos to a covert placebo (i.e., a placebo that entails deception surrounding its inert nature) administered in a physical reality setting (PR).

Employing an experimental heat-pain paradigm in healthy participants, we hypothesized that the placebo (a purported and “apparent” heat protective glove) would elicit placebo analgesia in the analgesia-expectation but not the control-expectation group in all three settings after the induction of analgesia-expectations in healthy participants; however, we expected the strongest placebo analgesia in PR (physical placebo, physical body), followed by AR (physical body, virtual placebo). We predicted that the VR condition (virtual body, virtual placebo) would also elicit placebo analgesia, but that this effect would be smaller compared to AR and PR, as weaker embodiment of the virtual (compared to the physical) body was predicted to reduce placebo responses. Furthermore, we expected that placebo analgesia in AR and VR would be influenced by the subjective level of embodiment, where stronger levels of embodiment are expected to predict stronger placebo responses. Lastly, we hypothesized that embodiment would predict pain perception, where participants would experience the least pain in PR, with AR again falling in the middle, and VR eliciting higher sensitivity to pain.

## 2. Methods

### 2.1 Ethics statement

The experiment was approved by the Cantonal Ethics Committee of Zurich (BASEC-Nr. 2017-02232) and all participants gave their written informed consent. The study was performed according to institutional ethics and national standards for the protection of human participants.

### 2.2 Participants

All participants were recruited through the University of Zurich “Fachverein Psychologie der Universität Zürich” mailing list. The study was introduced as an “investigation of pain perception in virtual and augmented reality”. Inclusion criteria were right handedness, proficient German language skills and being 18-65 years of age, whereas exclusion criteria were the presence of acute or chronic pain, pregnancy, drug or alcohol abuse, sensory abnormalities affecting thermal perception, a history of neurological or psychiatric disorders, or recent drug consumption. A sample size calculation using the software GPower [17] was conducted using an F-test for a repeated measures, within-between interaction MANOVA with a specified alpha set at 5% and power set at 80%, and an effect size of 0.55[9], which resulted in a total sample size of 49 participants. An initial group of 48 participants were recruited; however, in accordance with previous studies, participants with [14,64,65] low baseline pain threshold levels of *M* < 40°C, (n = 5) were excluded from further analyses. Therefore, additional five participants were recruited, with a final total of 48 participants used for analyses. Participants (19-62 years) were allocated to an analgesia-expectation (17 females, 7 males) (*M* = 24.71 years, *SD* = 4.7) or control-expectation (19 females, 5 males) (*M* = 27 years, *SD* = 9.14) group in a counterbalanced manner. The groups did not differ significantly in age [*t*(46) = 1.10, *p* = 0.28]. Participants received CHF 25 as monetary compensation for their participation, or one participant hour as fulfillment towards their academic requirements for the Bachelor of Science in Psychology at the University of Zurich. Although nausea and dizziness have previously been reported in experimental designs employing virtual reality, none of the participants in our sample reported any negative side effects.

### 2.3 Design

The experiment used a 2×3×2 mixed design with the between participants factor treatment group (analgesia-expectation, control-expectation) and the within factors setting (virtual, augmented, physical) and time (pre-intervention, post-intervention). Participants were assigned to either an analgesia-expectation or control-expectation group using an alternating quasi randomization allocation method, and order of settings was administered in a counterbalanced manner. Each participant completed an initial baseline pain threshold measurement, then completed three sessions of heat pain stimulation and subjective ratings for pain and embodiment, once in each setting, the order of which was also counterbalanced across participants (Fig. 1).

**Fig. 1.**
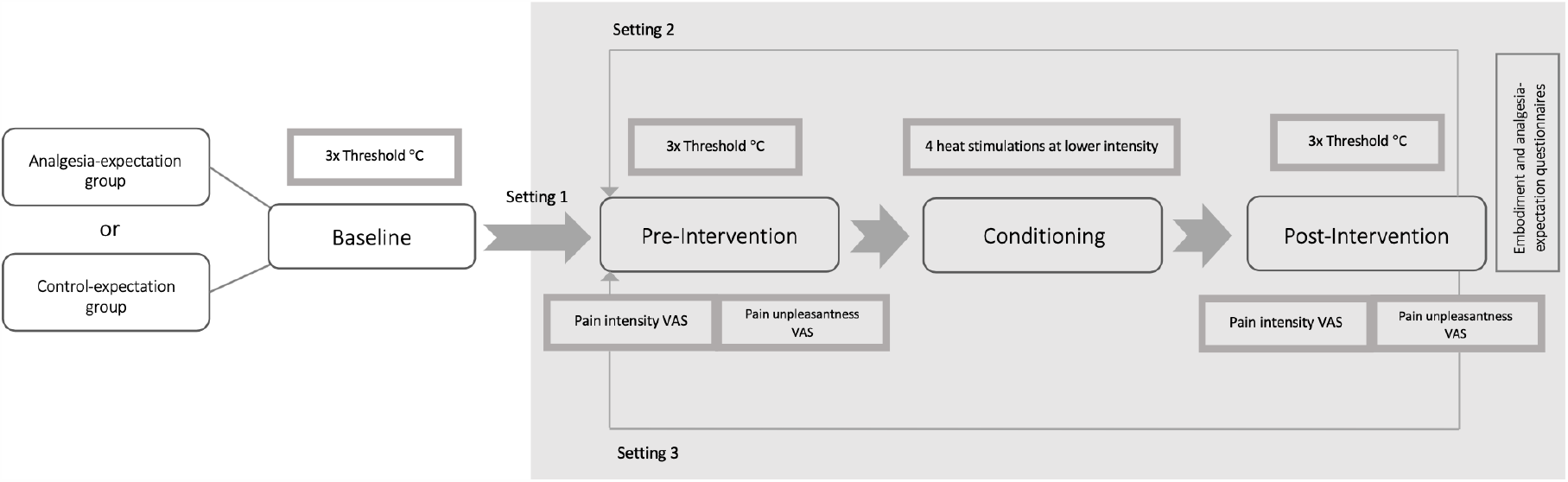
Experimental procedure. Participants were assigned to either an analgesia-expectation or control-expectation group and completed the experimental procedure in three settings (virtual reality setting, augmented reality setting, physical reality setting). Pain threshold, pain intensity, and pain unpleasantness were assessed pre-intervention and post-intervention.

### 2.4 Analgesia-expectation and control-expectation intervention

A glove reaching up to the elbow constituted the placebo (analgesia-expectation condition). Participants in the analgesia-expectation condition were informed that the aim of the study was to compare analgesia in three different settings (VR, AR, PR) in response to a heat-protective glove. They were informed that although they would not be wearing a physical glove in the virtual and augmented reality conditions, it would nevertheless be important that they report their pain experience as honestly and accurately as possible. When introducing the physical placebo to participants, it was presented deceptively as a novel, heat-protective glove, developed by the Swiss Institute of Technology, Zurich (ETHZ), that was made of nanotechnological materials designed to create micro sensory vibrations that deactivate C-fiber pain activation. They were instructed that they would see a virtual version of this glove in VR and AR. To further enhance the credibility of the scientific psychosocial context, the experimenter wore a white lab coat (for both analgesia-expectation and control-expectation groups) throughout the procedure [9,40]. At the end of each condition, participants in the analgesia-expectation group retrospectively rated analgesic efficacy of the heat protective glove, and how high their expectations were that the glove would protect them from the heat (Table 1).

**Table 1.** Analgesia-expectation questionnaire.

|  |  |
| --- | --- |
| 1. | How effective did you perceive the treatment to be? |
| 2. | How much did you expect that the glove would protect you from the heat? |

In contrast, participants in the control-expectation condition were informed that the aim of the experiment was to examine pain perception under different visual conditions, hence why they would either view the arm, or view a glove covering the arm.

Questions were only presented to participants in the analgesia-expectation treatment group.

### 2.5 Thermal stimulations

Heat pain measurement procedures were evoked using the TSA-II NeuroSensory Analyzer (Medoc, Ramat-Yishai, Israel). The thermode (30 mm × 30 mm) consisted of a metal plate that can be heated at various rates for different time periods, and was attached to the forearm. Prior to the administration of thermal stimuli, participants were familiarized with the device, thermode, and order of administration of the thermal stimuli. Baseline thresholds were administered to the left volar forearm at 1/3 of the distance from the wrist to the elbow. Participants were instructed to press the key to stop delivery of the stimulus when they reach the transition point of the thermal stimulus changing from “very hot” to “painful”. If the key was not pressed and to avoid physical injuries, thermal stimuli ceased at the predefined limit set at 52°C. Three baseline threshold stimuli were administered for calculation of the baseline threshold mean. Subsequently, the thermode was attached to the right lower forearm for commencement of the experimental procedure in the three settings. Prior to the start of each setting and to avoid sensory habituation from one setting to another, the thermode was shifted upwards, so that each setting corresponded with a new stimulation site (adapted from [9]), with a total of three differential sites used – one for each setting. Participants were always instructed to look at the arm being stimulated for each type of pain stimulation.

### 2.6 Settings

#### 2.6.1 Virtual reality

The immersive virtual environment was created with the Unity3D game engine (Unity version 2018.2.8). The corresponding visual stimuli were presented on an HTC Vive Pro (Vive™) head-mounted display with a resolution of 1440×1600 pixels per eye and a 110° diagonal field of view. Upper limb tracking was achieved using two HTC Vive Trackers (Vive™), one positioned on each of the participants’ lower arms, and the corresponding inverse kinematics were applied to move a virtual avatar in accordance with the participant’s movements, resulting in visuomotor correspondence. Steam VR (Steam®) was used to functionally combine the Vive kit with Unity.

The VR environment was designed to emulate the room conditions of the PR condition (Fig. 2A, 2B). Participants sat at a virtual desk that was colocalized to the physical desk. Two generic avatars (one male, one female) were used to match participants’ genders. The avatars were created in MakeHuman (www.makehumancommunity.org) and were viewed from a first-person perspective to ensure appropriate embodiment (Fig. 2C, 2D). In addition to the visuomotor correspondence, co-localization between the table in the PR condition and VR permitted additional visuotactile correspondence when participants would lay their arms on the table. A virtual thermode was attached to the virtual arm to emulate the physical conditions.

**Fig. 2.**
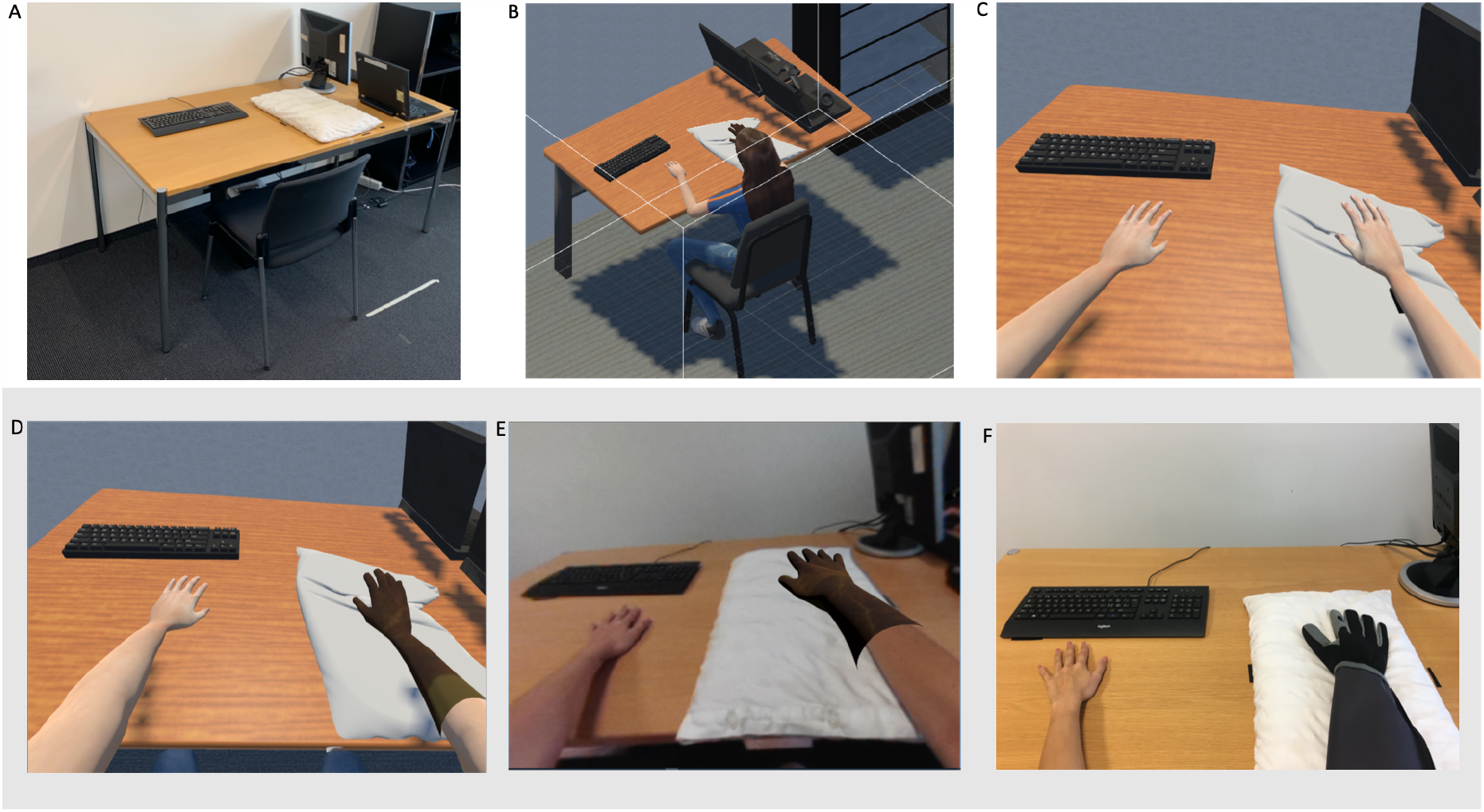
Experimental room in physical and virtual reality and placebo in virtual, augmented, and physical reality. The virtual experimental room (B) was designed to emulate the room in physical reality (A). A first-person perspective of the virtual avatar (C) together with synchronous visuomotor tracking was employed to induce the strongest sense of embodiment of the virtual avatar. In VR, the glove and body were presented as virtual objects (D), whereas only the glove was virtual in AR, while the own physical body was viewed through the headmounted display (E). In PR, participants viewed their own body and experienced a physical glove.

#### 2.6.2 Augmented reality

Participants saw the physical surroundings and their own physical body. The Vive Pro dual camera on the head mounted display was used to perform a real-time depth map and stereoscopic rendering of the environment using the Vive SRWorks Software Development Kit (Vive™). The same procedure used for VR was applied to track the participants’ movements and to present the visual stimulation. The placebo glove was virtually overlaid onto their physical arm (Fig. 2E).

#### 2.6.3 Physical reality

Participants did not wear the head-mounted display in this condition, so that the room, their body, as well as the placebo glove were experienced in a physical, “normal” PR condition (Fig. 2F). They only briefly wore the headset when answering questions pertaining to the experimental procedure in consistency with the other conditions.

### 2.7 Outcome assessments

#### 2.7.1 Pain thresholds

Each setting consisted of three pre-intervention and three post-intervention pain threshold stimulations, which started at 32°C baseline and increased at a rate of 1°C/s [23,32,34–36]. Participants were instructed to press the stop key on the keyboard as soon as they perceived the stimulation to change from “very hot” to “painful. Participants were instructed to always look at their arm (the one being stimulated) for each thermal stimulation.

#### 2.7.2 Pain ratings

Additionally, four pre- and four post-intervention individual pain stimulations were applied to measure pain intensity (“How painful was the stimulation?”) and pain unpleasantness (“How unpleasant was the stimulation?”) on a 0-1 visual analog scale (VAS) (values hidden from participants), ranging from “not painful at all” to “very painful”, and from “not unpleasant at all” to “very unpleasant”. Participants were told to differentiate between pain intensity and pain unpleasantness by providing an example that certain types of pain may be experienced as more intensely painful on a sensory level (e.g., knee pain), but could nevertheless be experienced as less unpleasant than another type of pain, which may be less intensely painful on a sensory level but perhaps more unpleasant on an affective level (e.g., headaches). These four individual pain stimulations were determined based on the individual’s calculated mean of the three baseline threshold stimulations and were set to [baseline threshold mean + 1°C], [baseline threshold mean + 1°C], [baseline threshold mean + 2°C], [baseline threshold mean + 2°C], which were always administered in this order. Temperature increased from a baseline temperature of 32°C at 10°C/s [9], maintained the individually calibrated temperature for three seconds, then decreased back down to baseline.

#### 2.7.3 Embodiment questionnaire

To measure the sense of embodiment experienced, participants completed an Embodiment Questionnaire (adapted from [9,51]) following the virtual and augmented reality sessions (Table 2). Questions were displayed within the headset visual space and were answered on a visual analog scale (VAS; values hidden from participants) from 0 (not at all) to 1 (very much) using headtracking to select and confirm answers. For the VR condition, the word “virtual arm” was used, whereas the word “seen arm” were employed in the AR and PR condition. The embodiment questionnaire in the PR condition was comprised only of questions emb1 and emb2. The questionnaire was adapted from commonly employed and validated [51] embodiment questionnaires in virtual bodily illusion experiments (e.g., [9]) (Table 2), and consisted of the following items: emb1. “How much did it feel like the pain you saw was caused by the stimulation on the virtual/seen arm?” (pain stimulation); emb2. “How strongly did it feel like the pain that you felt was caused at the same location on the virtual/seen arm?” (pain location); emb3. “How much did you feel like the virtual/seen arm is your own arm?” (ownership: the feeling that the seen arm is one’s own); emb4. “How strongly did it feel like the movements of the virtual/seen arms are your own movements?” (agency: the feeling that one is in control of one’s own movements); emb5. “How much did you feel like the virtual/seen body is a different person?” (lack of ownership: the feeling that the seen arm no longer constitutes one’s own body).

**Table 2.** Embodiment questionnaire.

|  |  |
| --- | --- |
| 1. | How much did it feel like the pain you saw was caused by the stimulation on the virtual/seen arm? |
| 2. | How strongly did it feel like the pain that you felt was caused at the same location on the virtual/seen arm? |
| 3. | How much did you feel like the virtual/seen arm is your own arm? |
| 4. | How strongly did it feel like the movements of the virtual/seen arm were your own movements? |
| 5. | How much did you feel like the virtual/seen body is a different person? |

In virtual and augmented reality, all five questions were presented to participants. In physical reality, only questions one and two were presented.

### 2.8 Experimental procedure

#### 2.8.1 Baseline

All experimental procedures were completed at the Department of Psychology at the University of Zurich. Participants were welcomed, seated at the desk, and informed on the experimental procedure, as well as on the TSA-II. The thermode was then applied to the lower left volar forearm and the three baseline pain thresholds were measured, the mean of which was then used for the four individual pain stimulation. Prior to beginning further experimental procedures, participants answered three short questions on mood (for additional mood measures, please see supplementary materials).

#### 2.8.2 Pre-intervention

Following baseline measurements, participants were administered to the first setting in the order determined by the Latin square counterbalancing scheme. Participants were instructed to move their arms side to side, as well as up and down – without rotating their wrists – to induce perceived embodiment of the virtual avatar, which moved in correspondence to their movements. To similarly assess perceived level of embodiment in AR and to further maintain the consistency across conditions, participants were instructed to briefly move their arms side to side and up and down in AR and PR. The thermode was then attached to the right lower volar forearm and participants were instructed to place their arms flat on the table. The right arm was placed on a white pillow for comfort and additional support, since the thermode was attached to the underside of the lower arm. Three pre-intervention pain thresholds were then measured for the first setting. After completion of the pain thresholds, four individual pain stimulations were administered. Following each stimulation, two questions appeared within the VR or AR environment, so that participants did not need to remove their headset. Each question was rated on a VAS from 0 (not painful/unpleasant at all) – 1 (very painful/unpleasant), though the numerical values remained hidden from participants. Participants saw a red dot that they were able to move using head movements, so that they could move the red dot along the VAS until it was located at the desired location for an answer. Participants could also correct their answers. Participants did not wear the head-mounted display in the PR condition, but were required to answer the questions through the headset to maintain consistency across conditions and avoid differential response styles.

#### 2.8.3 Expectation induction

After completion of the pre-intervention phase, the glove was applied to the right forearm. In the PR condition, the thermode remained under the glove. In the VR and AR conditions, the virtual glove appeared over the virtual (VR) or physical (AR) arm. In each condition, participants were told that the glove constituted a heat protective glove that will decrease pain from the heat stimulations. Effective induction of analgesia expectations can be achieved by lowering noxious stimulation levels (after an initial stimulus exposure), without informing participants about the real procedure [28,50,62]. In order to induce strong expectancy that the glove would decrease pain, participants were misleadingly informed that they would again receive the same four individual pain stimulations as during the pre-intervention phase; however, unbeknownst to participants, stimulation was surreptitiously reduced to [threshold - 1°C], [threshold - 1°C], [threshold - 2°C], and [threshold - 2°C].

#### 2.8.4 Post-intervention

The post-intervention phase immediately followed the expectation induction phase. Experimental procedures were identical to pre-intervention, where the temperature of the four individual pain stimulations was surreptitiously increased back to pre-intervention temperatures. Participants in the analgesia-expectation condition responded to an additional two questions surrounding the perceived retrospectively assessed analgesic efficacy (“How effective did you perceive the intervention to be?”) and expectations (“How much did you expect that the glove would protect you from the heat?”) of the placebo intervention (Table 1). All questions were presented in the HMD. Once all questions were completed, the experimental procedure started again at the pre-intervention phase for the next setting, until all three settings were completed. After the third setting condition, participants again rated their mood with identical items as to the baseline mood assessment [28]. Participants were then debriefed on the experimental procedures.

### 2.9 Statistical analyses

Data were analyzed with R studio Version 1.1.423 and were manually checked for assumptions of linear mixed model analyses (i.e., visual inspection of QQ plots and use of Shapiro Wilk test for normality) using the R *MASS* package [61]. Pre- to post-intervention difference scores were computed for thresholds, pain intensity ratings, and pain unpleasantness ratings (Δthresholds, Δpain intensity, Δpain unpleasantness). Independence of residuals and normal distributions were examined through inspections of QQ plots. Results from the QQ plots and the Shapiro-Wilk test (*p <* .05) showed that residuals were not normally distributed for Δthresholds, Δpain intensity, and Δpain unpleasantness ratings. Alpha was set at < .05, or 95% confidence intervals (CIs). Use of linear mixed model analysis was appropriate given the within-person dependence and the longitudinal structure, further allowing considering multiple observations across all participants for each task while adjusting for within-subject and within-group dependence. Since assumptions of normality do not apply to the current data, and the response variables do not fit a discrete distribution nor are binary, penalized quasilikelihood (PQL) was performed to adjust for non-normality.

The R-package brms [6], which is based on rstan [59] was used to calculate Bayesian ANOVAs and Bayesian multilevel models. Bayesian ANOVAs and multilevel models were calculated adopting the procedure described in Macauda and colleagues [31]. Bayesian procedures were used to provide posterior probability distributions for the estimated parameters, and non-informative priors were used for all parameters. The Hamilton Monte Carlo sampling algorithm was used to draw samples from each parameter’s posterior distribution [7]. Four independent Makov chain, each with 1000 warm-up samples, followed by an additional 1000 samples from the posterior distribution, were used to generate samples. The last 1000 samples of each Makov chain were saved for additional statistical inference. R-Hat statistics [7] were calculated to confirm that the samples for each chain converge on the same posterior distribution. Results show that all R-Hat statistics fell below 1.01, which demonstrated a low ratio of variance between he four chains to the variance within the chains. For the Bayesian multilevel models, we implemented the maximal random-effects structure justified by the experimental design [3], thus modeling by-participant random effects for each condition. The 95% Bayesian credible intervals (CI) for these posterior distributions constitute the probable range of the parameter considering the data and the model. If the CI does not contain zero, we can infer the existence of an effect.

## 3. Results

### 3.1 Baseline pain threshold

The Shapiro-Wilk test revealed that data were not normally distributed in the analgesia-expectation group (*W* = .93, *P* < .01); therefore, a Mann-Whitney test was used to calculate potential mean differences between analgesia-expectation and control-expectation groups for baseline pain threshold means. Baseline comparisons between groups indicated that mean baseline pain threshold did not differ significantly between the analgesia-expectation (*M* = 45.96) and control-expectation group (*M* = 45.97), *U* = 288, *P* = 1.00).

### 3.2 Pre-intervention to post-intervention differences

#### 3.2.1 Pain thresholds

Pain threshold increased for both the analgesia-expectation and control-expectation group between the pre- to post-intervention; however, the analgesia-expectation group seemed to display a markedly stronger increase in pain threshold from pre- to post-intervention. To prepare the data for a linear mixed model procedure, the difference between each of the three pre- and post-intervention pain threshold measures were calculated, so that a total of three scores (Δthreshold) remained per participant. Subsequently, a linear mixed model procedure including the random intercept but not slope examined potential differences between analgesia-expectation and control-expectation groups in pain thresholds with respect to setting (VR/AR/PR). Results from the linear mixed model analysis revealed no significant differences in pain thresholds between the three settings [*F*(2,382) = 0.25, *P* = 0.78]; however, there was a significant effect of group [*t*(46) = 5.016, *P* < .001, *SE* = .261] (Fig. 3), attributable to higher threshold differences from pre- to post-intervention for the analgesia-expectation group (*M* = 2.67) than the control-expectation group (*M* = 1.39).

**Fig. 3.**
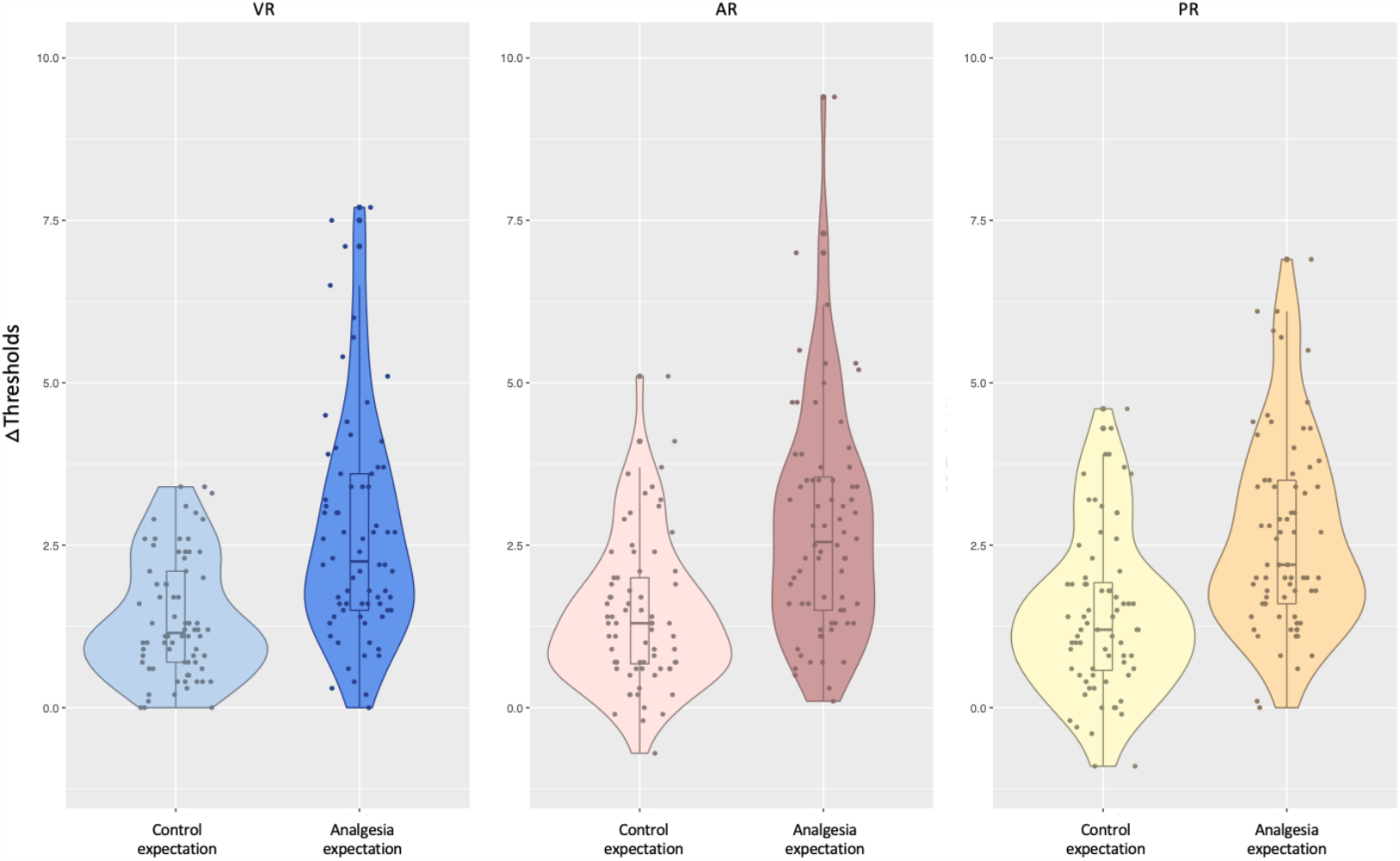
Group median, interquartile ranges, and individual values (difference between the average of the individual three pre-intervention and three post-intervention thresholds) for pre-intervention to post-intervention differences in thresholds in analgesia-expectation (dark colors) and control-expectation groups (light colors) in the three settings. Participants in the analgesia-expectation group demonstrated larger pre-intervention to post-intervention increases in thresholds than participants in the control-expectation group across VR (blue), AR (pink) and PR (yellow) settings.

With the implementation of a Bayesian multilevel model, we examined the effect of setting (VR vs. AR vs. PR) and the effect of group (analgesia-expectation vs. control-expectation) on pre- to post-intervention threshold differences. The Bayesian multilevel model confirmed no effect of setting (i.e., CI-MPE crossing the zero), therefore confirming the null hypothesis, but did confirm a significant effect of group (MPE = 1.31, 95% CI = [0.75, 1.86]), indicating significant higher thresholds in the analgesia-expectation than the control-expectation group.

#### 3.2.2 Pain intensity

Data for pain intensity were similarly prepared for a linear mixed model analysis by computing the difference score between pre- to post-intervention ratings (Δpain intensity), with again three remaining Δpain intensity scores representing the difference from pre- to post-intervention for each participant. Linear mixed model analysis again examined potential differences between analgesia-expectation and control-expectation groups for pain intensity ratings with respect to setting (VR/AR/PR) experienced. Results from the linear mixed model analysis again revealed no significant differences in pain intensity ratings for the three settings [*F*(2,382) = 0.02, *P* = .98], but there again was a significant effect of group [*t*(46) = −3.78, *P* < .001, *SE* = .03)] (Fig. 4A). Participants in the analgesia-expectation group (*M* = −0.18) reported a greater decrease in pain intensity from pre- to post-intervention than participants in the control-expectation group (*M* = −0.03).

**Fig. 4.**
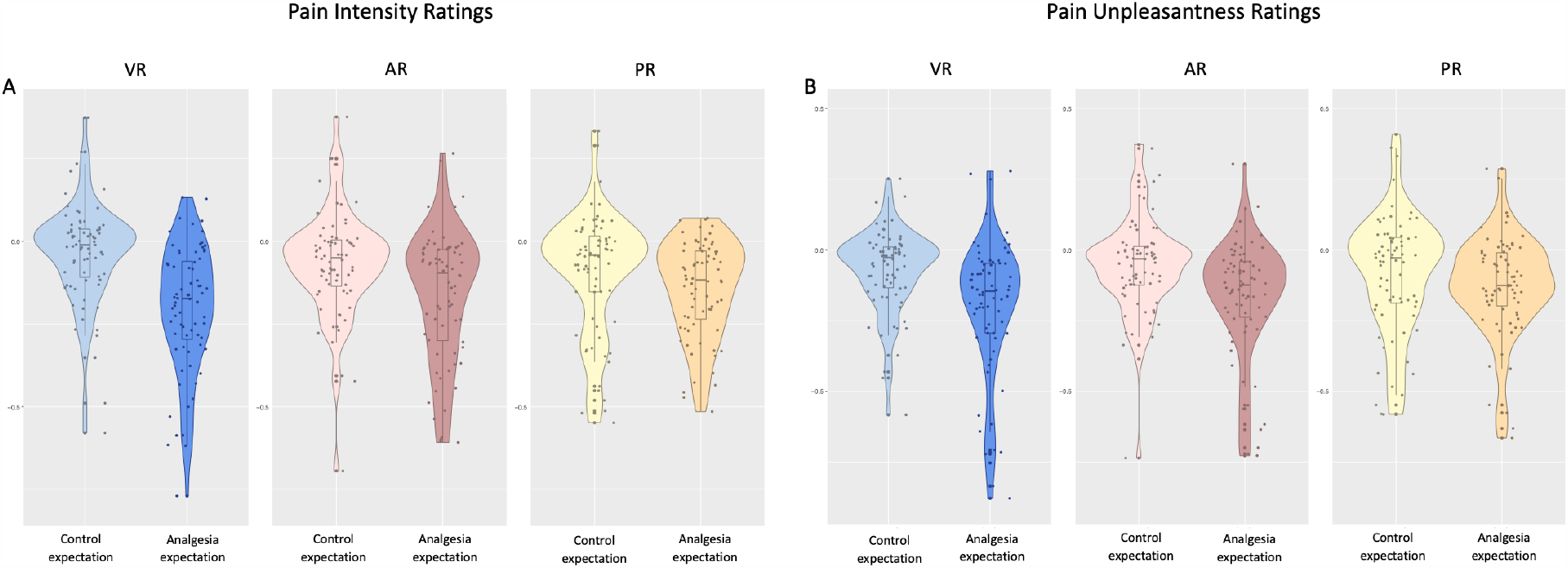
Group median, interquartile ranges, and individual values (difference between the average of the individual four pre-intervention and four-post intervention ratings) for pre-intervention to post-intervention differences in sensory pain ratings (A) and pain unpleasantness ratings (B) in analgesia-expectation (dark colors) and control-expectation (light colors) groups in the three settings. Participants in the analgesia-expectation group reported larger decreases in pre-intervention to post-intervention pain intensity ratings in VR (blue), AR (pink) and PR (yellow) than participants in the control-expectation group for both pain intensity (A) and pain unpleasantness (B) ratings.

The Bayesian multilevel model showed no effect of setting (i.e., CI-MPE crossing the zero) but again a significant effect of group (MPE = −0.11, 95% CI = [−0.16, −0.05]), indicating significant lower pain intensity ratings in the analgesia-expectation than the control-expectation group.

#### 3.2.3 Pain unpleasantness

Data for pain unpleasantness ratings again were prepared for a linear mixed model analysis by computing the difference between the pre- to post-intervention score (Δpain unpleasantness), with a resulting total of three Δpain unpleasantness scores for each participant. The linear mixed model analysis again examined potential differences between groups (analgesia-expectation/control-expectation) and the three settings (VR/AR/PR). Results here as well demonstrated that there were no significant differences in the setting experienced [*F*(2,382) = 1.25, *P* = 0.29], but there was a significant effect of group [*t*(46) = −2.98, *P* < .01, *SE* = 0.03)] (Fig. 4B). Participants in the analgesia-expectation group (M = −0.18) showed greater decreases in pre- to post-intervention pain unpleasantness ratings than participants in the control-expectation group (M = −0.07).

Using a Bayesian multilevel model, we examined the effect of setting (VR vs. AR vs. PR) and the effect of group (analgesia-expectation vs. control-expectation) on pre- to post-intervention threshold differences. The Bayesian multilevel model showed no effect of setting (i.e., CI-MPE crossing the zero) but a significant effect of group (MPE = −0.10, 95% CI = [−0.16, −0.03]), indicating significant lower pain unpleasantness ratings in the analgesia-expectation than the control-expectation group.

### 3.3 Pre-intervention pain perception

#### 3.3.1 Pain thresholds

We used a linear mixed model procedure to examine whether pain thresholds differed according to setting (VR/AR/PR) experienced. Results revealed no significant differences in pre-intervention pain thresholds between the three settings [*F*(2,382) = 0.94, *P* = 0.39].

Using a Bayesian multilevel model, we examined the effect of setting (VR vs. AR vs. PR) and the effect of group (analgesia-expectation vs. control-expectation) on pre-intervention pain thresholds. The Bayesian multilevel model showed no significant effect of setting and no significant effect of group (i.e., CI-MPE crosses zero).

#### 3.3.2 Pain intensity

We again used a linear mixed model procedure to examine whether pain intensity differed according to setting (VR/AR/PR) experienced. Results revealed a significant difference in pre-intervention pain intensity ratings between the three settings [*F*(2,670) = 3.46, *P* < .05]. While there was no significant difference between PR and AR [*t*(670) = 0.58, *P* = .56, *SE* = .02)], there was a trend in pain intensity between VR and AR [*t*(670) = −1.93, *P* = .054, *SE* = .02)], and a significant difference between VR and PR [*t*(670) = 2.51, *P* < .05, *SE* = .02)] (Fig. 5). Participants reported lower pain intensity ratings for VR (*M* = 0.37) than in PR (*M* = 0.43).

**Fig. 5.**
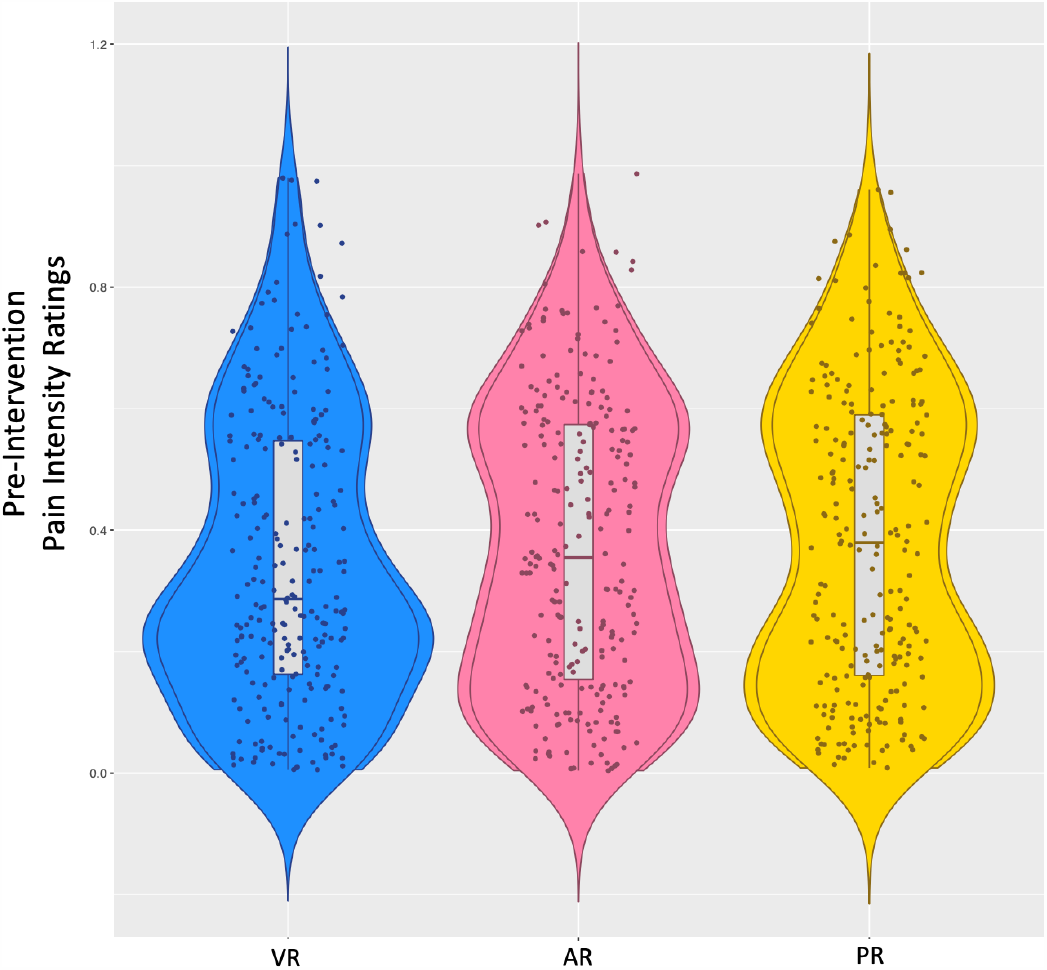
Group median and individual values (mean of the four pre-intervention pain ratings) for pre-intervention pain intensity ratings in VR, AR, and PR. Participants reported significantly lower pain ratings during pre-intervention in VR (blue) than in AR (pink) or PR (yellow).

Using a Bayesian multilevel model, we examined the effect of setting (VR vs. AR vs. PR) and the effect of group (analgesia-expectation vs. control-expectation) on pre-intervention pain intensity ratings. The Bayesian multilevel model showed a significant effect of setting between PR and VR (MPE = 0.038, 95% CI = [0.01, 0.07]), indicating higher pain intensity ratings in PR compared to VR, but no significant differences between AR and PR, and between AR and VR (i.e., CI-MPE crosses zero).

#### 3.3.3 Pain unpleasantness

We again used a linear mixed model procedure to examine whether pain unpleasantness differed according to setting (VR/AR/PR) experienced. Results revealed that differences in pre-intervention pain unpleasantness ratings between the three settings were just barely not significant [*F*(2,670) = 2.99, *P* = .0509).

Using a Bayesian multilevel model, we examined the effect of setting (VR vs. AR vs. PR) and the effect of group (analgesia-expectation vs. control-expectation) on pre-intervention pain unpleasantness. The Bayesian multilevel model showed no significant effect of setting and no significant effect of group (i.e., CI-MPE crosses zero).

### 3.4 Embodiment

#### 3.4.1 Embodiment comparison: Analgesia-expectation and control-expectation

Embodiment scores (means, ranges, and standard deviations) are listed in Table 3 for the VR condition and in Table 4 for the AR condition. To assess perceived embodiment between analgesia-expectation and control-expectation groups for VR and AR, a multivariate analysis of variance (MANOVA) was conducted on the five embodiment questions for VR and AR. The analysis did not reveal a significant effect of group in the VR [Pillai’s *V* = 0.13, *F*(1,46) = 1.27, *P* > .05] or AR [Pillai’s *V* = 0.17, *F*(1,46) = 1.73, *P* > .05] condition, demonstrating that the groups did not exhibit differences in subjective embodiment.

**Table 3.** Means, ranges, and standard deviations (SD) for subscales of the embodiment questionnaire in VR.

|  | Pain stimulation | Pain location | Ownership | Agency | Lack of ownership | Composite |
| --- | --- | --- | --- | --- | --- | --- |
| Valid | 24 | 24 | 24 | 24 | 24 | 24 |
| Mean | 0.615 | 0.727 | 0.599 | 0.722 | 0.329 | 3.663 |
| SD | 0.235 | 0.231 | 0.294 | 0.198 | 0.215 | 0.709 |
| Minimum | 0.241 | 0.163 | 0.064 | 0.134 | 0.027 | 2.296 |
| Maximum | 0.979 | 0.985 | 0.977 | 0.987 | 0.907 | 4.869 |

**Table 4.** Means, ranges, and standard deviations (SD) for subscales of the embodiment questionnaire in AR.

|  | Pain stimulation | Pain location | Ownership | Agency | Lack of ownership | Composite |
| --- | --- | --- | --- | --- | --- | --- |
| Valid | 24 | 24 | 24 | 24 | 24 | 24 |
| Mean | 0.727 | 0.805 | 0.766 | 0.807 | 0.192 | 4.104 |
| SD | 0.215 | 0.153 | 0.236 | 0.238 | 0.221 | 0.662 |
| Minimum | 0.168 | 0.487 | 0.045 | 0.024 | 0.007 | 2.696 |
| Maximum | 0.997 | 0.996 | 0.996 | 0.997 | 0.958 | 4.983 |

#### 3.4.2 Embodiment comparison: VR and AR

To compare embodiment ratings in VR and AR, paired samples t-tests between VR and AR in both analgesia-expectation and control-expectation groups for composite embodiment scores and each of the embodiment questions were conducted, which revealed significant differences in perceived embodiment between the two settings. For composite embodiment scores, participants felt significantly more embodied in the AR (0.75 ± 0.17) than in the VR (0.62 ± 0.17) condition [*t*(47) = −6.34, *P* < .001]. In the AR condition (0.69 ± 0.21), participants perceived more strongly that the pain that they saw was caused by the stimulation on the virtual/seen arm than in the VR condition (0.58 ± 0.25) [emb1: *t*(47) = −4.32, *P* < .001]. Similarly, participants also felt more strongly that the pain that they felt was caused at the same location on the virtual/seen arm in the AR condition (0.74 ± 0.18) than in the VR condition (0.66 ± 0.52) [emb2: *t*(47) = −2.95, *P* < .01]. The AR condition (0.75 ± 0.21) also elicited more ownership over the arm than in VR (0.54 ± 0.26) [emb3: *t*(47) = −6.14, *P* < .001]. The AR condition (0.76 ± 0.26) also elicited stronger feelings of agency over the arm than in VR (0.67 ± 0.19) [emb4: *t*(47) = −2.49, *P* < .01]. Lastly, the feeling that the virtual/seen body was a different person was weaker in AR (0.21 ± 0.22) than in VR (0.36 ± 0.20) [emb5: *t*(47) = 5.68, *P* < .001] (Fig. 6).

**Fig. 6.**
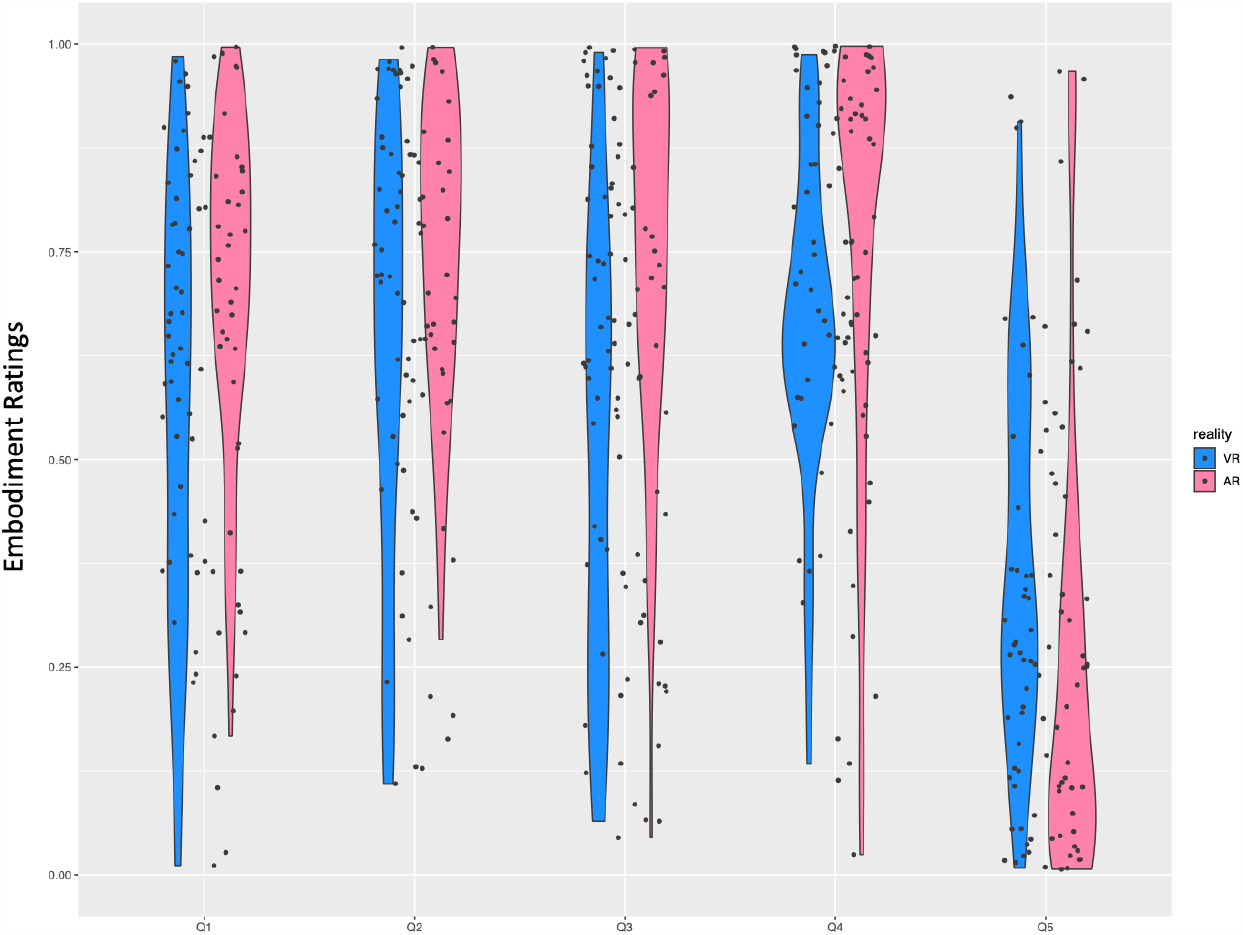
Individual values for perceived embodiment ratings in VR (blue) and AR (pink) of the embodiment questionnaire. Participants in the AR setting felt more strongly that they experienced the pain more on their own seen arm than the virtual arm (Q1), that it was caused at the same location (Q2), that the seen arm was their own arm (Q3), that the movements were their own movements (Q4), and felt less strongly that the seen arm was a different person (Q5) than in the VR setting.

#### 3.4.3 Embodiment and placebo analgesia

Embodiment was examined as a predictor of placebo responses, which included retrospectively assessed analgesic efficacy, thresholds, pain intensity, and pain unpleasantness differences in the analgesia-expectation group only. Embodiment subscales were collapsed into a single composite embodiment mean score for each participant, where lack of ownership (emb5: “How much did you feel like the virtual body is a different person?”) was reverse coded. A multivariate linear regression was calculated to predict the effects of VR and AR embodiment and retroactively assessed analgesia expectation on Δthresholds, Δpain intensity, and Δpain unpleasantness.

In the VR condition, a significant regression equation was found [F(2,20) = 15.66, *P* < .001], with an adjusted *R*^*2*^ of 0.57. Participants’ predicted retrospectively assessed analgesic efficacy is equal to −0.29 – 0.77 (retrospectively assessed analgesia expectation), and predicted retrospectively assessed analgesic efficacy is equal to −0.29 – 0.66 (embodiment), where data for retrospectively assessed analgesic efficacy, retrospectively assessed analgesia expectation, and embodiment range from 0-1 on a continuous scale. Therefore, both retrospectively assessed analgesia expectation and embodiment constituted significant predictors of retrospectively assessed analgesic efficacy (Fig. 7). No significant regression equation was found for retrospectively assessed analgesia expectation as a predictor.

**Fig. 7.**
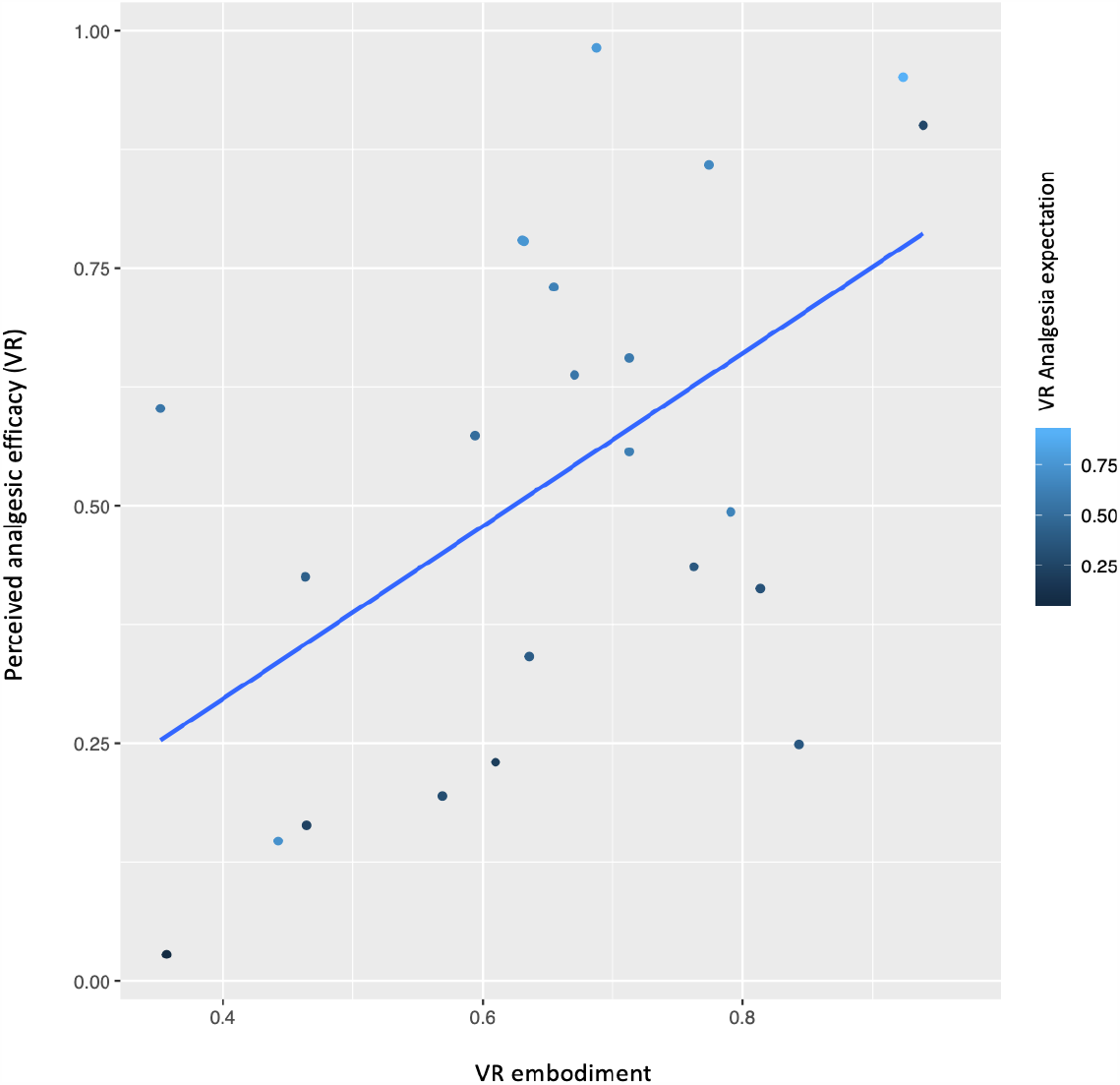
Standardized values of residuals and average predicted analgesic efficacy of placebo analgesia by strength of embodiment in VR. Retrospectively assessed analgesia expectation and embodiment significantly predicted retrospectively assessed analgesic efficacy.

In the AR condition, embodiment did not significantly predict placebo analgesia, but a significant regression equation was found for retrospectively assessed analgesia expectation, where retrospectively assessed analgesia expectation predicted retrospectively assessed analgesic efficacy [F(2,20) = 6.71, *P* < .01], with an adjusted *R*^*2*^ of .34. Participants’ predicted retrospectively assessed analgesic efficacy is equal to 0.00 – 0.74 (retrospectively assessed analgesia expectation). Retrospectively assessed analgesia expectation further predicted Δthresholds [F(2,20) = 2.65, *P* < .05], with an adjusted *R*^*2*^ of 0.13. Participants’ Δthresholds is equal to 0.98 – 3.10 (retrospectively assessed analgesia expectation), where retrospectively assessed analgesia expectation was coded on a VAS (0-1) and thresholds were measured in C. Additionally, there was a significant regression equation found for Δpain unpleasantness [F(2,20) = 7.77, *P* < .01], with an adjusted *R*^*2*^ of 0.38. Participants’ Δpain unpleasantness is equal to −0.15 – (−0.45) (retrospectively assessed analgesia expectation), where both pain unpleasantness and retrospectively assessed analgesic efficacy range from 0-1 on a VAS. While it did not reach significance, there was a trend that retrospectively assessed analgesia expectation similarly predicts Δpain intensity (*P* = .08). These results suggest that retrospectively assessed analgesia expectation was a significant predictor of pre- to post-intervention increases in thresholds, and decreases in pain unpleasantness.

In the PR condition, participants were asked the first two embodiment questions only, pertaining to pain stimulation and pain location. Multivariate regression analysis revealed no significant regression equations for retrospectively assessed analgesia expectation or embodiment (pain stimulation and pain location).

### 3.5 Embodiment and pain perception

Embodiment was examined as a predictor of pain perception in the pre-intervention phase, which included pre-intervention thresholds, pain intensity, and pain unpleasantness ratings. A multivariate linear regression was calculated to predict the effects of VR and AR embodiment on pre-intervention thresholds, pain intensity, and pain unpleasantness. Embodiment subscales were again collapsed into a single composite embodiment mean score for each participant, where lack of ownership (emb5: “How much did you feel like the virtual body is a different person?”) was reverse coded. However, no significant regression equations were found for embodiment in VR on pre-intervention measures of thresholds [F(1,46) = 0.70, *P* = .41], with an adjusted *R*^*2*^ of −0.01, pain intensity [F(1,46) = 2.86, *P* = .10], with an adjusted *R*^*2*^ of 0.04, or pain unpleasantness [F(1,46) = 2.00, *P* = .17], with an adjusted *R*^*2*^ of 0.02, nor in AR on pre-intervention measures of thresholds [F(1,46) = 1.14, *P* = .29], with an adjusted *R*^*2*^ of 0.003, pain intensity [F(1,46) = 2.06, *P* = .16], with an adjusted *R*^*2*^ of 0.24, or pain unpleasantness [F(1,46) = 1.46, *P* = .23], with an adjusted *R*^*2*^ of 0.01.

### 3.6 Additional analyses

For additional correlational analyses of mood on outcome measures, please see the supplementary materials.

## 4. Discussion

This study reports the first experimental evidence for successful expectancy-induced virtual placebo analgesia working as efficiently when administered to an embodied virtual instead of a physical body in healthy participants. Participants in the analgesia-expectation group displayed larger increases in pain threshold, as well as greater reductions in pain intensity and pain unpleasantness ratings from pre- to post-intervention compared to the control group. Contrary to our expectations, we found no differences in analgesic strength between an open label, virtual placebo compared to its covert counterpart in physical form. As such, the present study suggests that a placebo does not need to necessarily be applied in physical and covert form, or even to a physical body, to still successfully increase experimental pain thresholds and reduce pain ratings. Unlike predicted, this effect was overall not modulated by the sense of embodiment, despite results that embodiment was stronger in AR than VR. However, as a partial confirmation of our hypotheses, our results evince that embodiment predicts some aspects of placebo analgesia in VR, but not in AR. Furthermore, baseline pain intensity ratings were significantly lower in VR than the other settings, yet this effect was not confirmed in the pain threshold and pain unpleasantness measures, nor did embodiment predict pain perception.

### 4.1 No significant differences in analgesic efficacy between an open label and physical placebo

We hypothesized that placebo analgesia would be strongest in PR, followed by AR, and then VR. However, our results suggest no differences between a placebo administered in VR, AR and PR, despite the covert nature of the placebo (PR) that contrasted the open label placebos (OLP) in VR and AR. For a long time, it has been presumed that placebo substances can produce therapeutic benefits only if patients were unaware they had received a sham treatment or medication. Recently, however, the notion that placebo analgesia necessitates deception or concealment has been challenged, as administering OLP with rationale can be as efficacious as providing covert placebos [8,10,27,29].

While direct evidence of computational models in placebo analgesia are still quite preliminary, prediction error processing seems to fit into the mechanistic neuropsychological explanatory models of placebo responses [5,27] and pain modulation [63]. Prediction error processing could correct the neurological, cognitive, and bodily dissonance during the contradictory messages experienced when administered OLP (i.e., the placebo could help or the placebo cannot work), resulting in nonconscious inferences that disturb central sensitization [27,45]. Bayesian inferencing becomes particularly pertinent in the presence of a high degree of uncertainty and when priors (neutrally encoded probability distributions that are contrasted against posterior sensory evidence [55]) are ambiguous, such as during administration of OLP [27], which could have been further increased by the novel, virtual nature of the placebos in this study. Perceived embodiment of a virtual body integrated in a scientific setting with plausible rationales, experimental rituals and psychosocial cues, such as a white lab coat and attributes or characteristics of the experimenter (e.g., competent demeanor) [25], could induce expectancy-related analgesia involving a top-down activation of endogenous analgesic activity via the descending pain modulatory system [4,13,15].

Our results extend existing literature on placebo analgesia by demonstrating that the efficacy of OLP extend beyond physical boundaries to virtual settings (and bodies) as well. Future studies may consider experimenting with different types of virtual placebos (e.g., a virtual analgesic cream) or examining psychophysiological interactions with placebo analgesia response inhibitors, such as naloxone [15] or non-invasive neurostimulation [28].

### 4.2 Placebo analgesia and embodiment

We hypothesized that placebo analgesia in AR and VR would be influenced by the subjective level of embodiment, where stronger levels of embodiment would predict stronger placebo responses [9]. Contrary to our expectations, placebo analgesia did not differ across settings, despite varying levels of embodiment of the physical versus virtual body. Interestingly, however, embodiment predicted retrospectively assessed analgesic efficacy only in VR, where generally lower embodiment scores were found compared to AR.

While this effect will necessitate further investigation by including conditions aimed at reducing the sense of embodiment (e.g., asynchronous multisensory stimulation) [51], we speculate that an embodied approach could provide a plausible explanation for our results. Bodily perception epitomizes a foundational basis of our self: even if higher order features of consciousness are reduced, a basic sense of self embedded in a body is retained [21], emphasizing that all experience occurs from within this embodied frame of reference [19,20,22]. Embodied approaches to computational self-models postulate that domain specific priors surrounding body awareness assume a precedence status in the cortical hierarchy due to the organism’s natural impetus in maintaining homeostasis [1]. Whereas our bodily sensations generally go unnoticed in our daily lives, novel situations, such as embodiment of a virtual body, can increase the salience of attention to such bodily cues. Changes in stimulus intensity of exogenous attentional effects and bottom-up visual, tactile, and proprioceptive information could therefore increase the weight of bottom-up cues surrounding the (virtual) body, while decreasing the comparative salience of treatment context induced top-down expectations surrounding the placebo procedure (i.e., psychosocial cues and rituals). In the presence of VR-specific body-related novelty or uncertainty, comparatively greater relevance might be placed on embodiment compared to other contextual factors. While embodiment did not predict pre- to post-intervention differences for threshold, pain intensity, or pain unpleasantness, the purported precedence of embodiment may constitute a prerequisite in experiencing the perceived efficacy of placebo analgesia in VR.

In AR, however, participants saw a video-based depiction of their own body through the head-mounted display, thereby lowering the level of attention and uncertainty surrounding the body and thus maintaining low body-related prediction error. We suggest that these differences may redirect attention and processing to higher cortical levels, specifically those related to expectations. In the AR condition, increased weight may be given to expectations partially due to the comparative smaller relevance of body-specific processing, but also due to the nature of the placebo. While the open label nature of the placebo applied to both the VR and AR conditions, embodying the own body in AR could have constituted a driving factor in shifting a greater weight to conscious expectancy in producing placebo responses.

### 4.3 Pain perception and embodiment

The study of the relationship between pain and embodiment has a long tradition. The sense of embodiment has been suggested as a modulator of pain perception [33,52], yet empirical results are inconsistent [39,44]. In the current study, embodiment was successfully modulated by the different settings as measured by a questionnaire. We hypothesized that embodiment would predict pain perception, where participants would experience the least pain in PR, with AR falling in the middle, and VR eliciting higher sensitivity to pain. Contrary to our expectations, participants reported significantly lower pain ratings in VR than in PR; however, no differences in pain threshold and unpleasantness were found, suggesting that viewing an embodied virtual arm may evince similar analgesic effects to viewing the own body. While viewing the limb encompasses analgesic qualities [30], simply viewing someone else’s or an unembodied (virtual) limb does not seem to significantly affect pain, suggesting that embodiment of the artificial or virtual body might constitute a mediating factor [35]. However, our results do not evince embodiment as a significant predictor of pre-intervention threshold, pain intensity, or pain unpleasantness when examined in VR and AR. These seemingly contradictory results could be explicated by our study design aimed at maximizing perceived embodiment: although participants generally felt more embodied of their own physical body in AR, synchronous visuomotor coherence generally ensures illusory ownership of the virtual body. However, it should be noted that attention can strongly influence pain perception [26,46], and should therefore also be considered as a potential modulating factor in these results. Immersive VR was an interactive experience in a novel setting, eliciting a feeling of presence that could decrease pain perception through means of distraction [24]; therefore, future studies may consider disentangling these factors. Nevertheless, while previous studies have demonstrated that ownership of the virtual body seems to constitute a crucial role in experiencing analgesic effects [23,33,34,52], the current study suggests that the alteration of illusory embodiment is sufficient to elicit analgesic effects, independent of perceived strength.

### 4.4 Conclusions and outlook

Virtual reality provides a high degree of sensory control, permitting the construction of novel virtual environments that can influence and systematically test context-mediated effects in novel ways. Our findings suggest no differences in analgesic efficacy between a virtual, OLP and a covert, physical placebo, and that a placebo administered to an embodied virtual body could influence treatment expectations akin to a placebo administered to the physical body. This is even more promising, as our data suggest that the perception of pain intensity is decreased in VR pre-intervention. Based on models of predictive coding, we propose that the foundational sense of embodiment of a virtual body assumes a central role in the contextual processing of placebo analgesia in VR, which could pave the way for further, more elaborate investigations of how embodiment, and modulations thereof, influence the perception of pain. Future research would benefit from examining the boundaries of these effects by including variations in the strength of embodiment (e.g., using asynchronous stimulation) or introducing modulations to the virtual body. Although the current study focused on experimentally induced acute pain, the results could nevertheless serve as a promising foundational framework for non-invasive, non-pharmacological approaches focused on employing embodied virtual reality as a potential tool for pain management. Though additional research will be needed for chronic pain, introducing virtual placebos could potentially act synergistically with other (virtual) bodily modifications that target body perception disturbances, which have already demonstrated considerable success in reducing pain in several pain types [38,41,42].

JTH, BL and MRL were funded by the Swiss National Science Foundation (grant number: P00P1_170511), PK was supported by the crowdfunding project “Heilen mit Hirn”. There are no conflicts of interest. We would like to thank Peter Brugger for his contribution and insights to the experimental design and manuscript, and Gianluca Saetta for his support in Bayesian statistics.

## Supporting information

Supplemental Analyses Mood

Supplemental Tables Mood

## Notes

### Competing Interest Statement

The authors have declared no competing interest.

