## Supplemental Analyses Mood for "Real Bodies Not Required? Placebo Analgesia and Pain Perception in Immersive Virtual and Augmented Reality"

**Table 1-SM.**

Spearman’s Rank correlations between embodiment and placebo analgesia in VR in the analgesia-expectation group only. Spearman’s Rank correlations were performed to adjust for nonparametric data, and adjusted for multiple comparisons (new significance value = *P* < .004). ΔThreshold (Threshold Diff), Δpain intensity (Pain Intensity Diff), and Δpain unpleasantness (Pain Unpleasantness Diff) were correlated with embodiment questions (VR and AR: five embodiment questions Emb Q1 – Emb Q5; PR: Emb Q1 – Emb Q2), as well as with analgesia expectation and perceived placebo efficacy responses. Better mood post experiment negatively correlated with pre to post intervention pain intensity [*r_s_*(12) = -0.68, *P* < .001] and pain unpleasantness ratings [*r_s_*(12) = -0.64, *P* < .01], meaning that higher ratings for good mood correlated to greater decreases from pre-intervention to post-intervention in pain intensity and unpleasantness. Furthermore, calmer mood post experiment correlated positively with analgesia expectation [*r_s_*(12) = 0.62, *P* < .01]. No other significant correlations were found (all comparisons *P* > .007, *r*(12) > -0.55).

**Table 2-SM.**

Spearman’s Rank correlations between embodiment and placebo analgesia in AR in the analgesia-expectation group only. Spearman’s Rank correlations were performed to adjust for nonparametric data, and adjusted for multiple comparisons (new significance value = *P* < .004). ΔThreshold (Threshold Diff), Δpain intensity (Pain Intensity Diff), and Δpain unpleasantness (Pain Unpleasantness Diff) were correlated with embodiment questions (VR and AR: five embodiment questions Emb Q1 – Emb Q5; PR: Emb Q1 – Emb Q2), as well as with analgesia expectation and perceived placebo efficacy responses. After correcting for multiple comparisons, no correlations in the AR condition maintained significance (all comparisons *P* > .007, *r*(12) > -0.54).

**Table 3-SM.**

Spearman’s Rank correlations between embodiment and placebo analgesia in PR in the analgesia-expectation group only. Spearman’s Rank correlations were performed to adjust for nonparametric data, and adjusted for multiple comparisons (new significance value = *P* < .004). ΔThreshold (Threshold Diff), Δpain intensity (Pain Intensity Diff), and Δpain unpleasantness (Pain Unpleasantness Diff) were correlated with embodiment questions (VR and AR: five embodiment questions Emb Q1 – Emb Q5; PR: Emb Q1 – Emb Q2), as well as with analgesia expectation and perceived placebo efficacy responses. After correcting for multiple comparisons, no correlations in the PR condition maintained significance (all comparisons *P* > .007, *r*(12) > -0.54).

**Table 4-SM.**

Spearman’s Rank correlations between mood and placebo analgesia in VR in the control-expectation group only. Spearman’s Rank correlations were performed to adjust for nonparametric data, and adjusted for multiple comparisons (new significance value = *P* < .004). ΔThreshold (Threshold Diff), Δpain intensity (Pain Intensity Diff), and Δpain unpleasantness (Pain Unpleasantness Diff) were correlated with mood questions (GB: good-bad; AT: awake-tired; CN: calm-nervous). Following corrections for multiple comparisons, no correlations maintained statistical significance (all comparisons *P* > .03, *r*(10) > -0.44).

**Table 5-SM.**

Spearman’s Rank correlations between mood and placebo analgesia in AR in the control-expectation group only. Spearman’s Rank correlations were performed to adjust for nonparametric data, and adjusted for multiple comparisons (new significance value = *P* < .004). ΔThreshold (Threshold Diff), Δpain intensity (Pain Intensity Diff), and Δpain unpleasantness (Pain Unpleasantness Diff) were correlated with mood questions (GB: good-bad; AT: awake-tired; CN: calm-nervous). Calmer mood post experiment correlated significantly with pre to post decreases in pain intensity ratings [*r_s_*(10) = -0.78, *P* < .001], where participants who had higher ratings for calmness following the experiment also reported greater decreases in pain intensity from pre-intervention to post-intervention. Following corrections for multiple comparisons, no correlations maintained statistical significance (all comparisons *P* > .03, *r*(10) > -0.46).

**Table 6-SM.**

Spearman’s Rank correlations between mood and placebo analgesia in AR in the control-expectation group only. Spearman’s Rank correlations were performed to adjust for nonparametric data, and adjusted for multiple comparisons (new significance value = *P* < .004). ΔThreshold (Threshold Diff), Δpain intensity (Pain Intensity Diff), and Δpain unpleasantness (Pain Unpleasantness Diff) were correlated with mood questions (GB: good-bad; AT: awake-tired; CN: calm-nervous). A significant positive correlation was found between feeling more awake post experiment and Δthreshold [*r_s_*(10) = 0.70, *P* < .001], where participants who reported they felt more awake after completion of the experiment also demonstrated greater pre-intervention to post-intervention increases in thresholds. Following corrections for multiple comparisons, no correlations maintained statistical significance (all comparisons *P* > .02, *r*(10) > -0.48).
