## Supplemental Tables Mood for "Real Bodies Not Required? Placebo Analgesia and Pain Perception in Immersive Virtual and Augmented Reality"

Table 1 – Supplementary Materials.

| **Spearman Correlations: Mood and Placebo Analgesia in VR** | | | | | | | | |
| --- | --- | --- | --- | --- | --- | --- | --- | --- |
|  | |  | |  | | **Spearman’s rho** | | **p** |
| Threshold diff |  | - |  | Mood pre GB |  | -0.293 |  | 0.164 |
| Threshold diff |  | - |  | Mood pre AT |  | 0.104 |  | 0.627 |
| Threshold diff |  | - |  | Mood pre CN |  | -0.028 |  | 0.897 |
| Threshold diff |  | - |  | Mood post GB |  | 0.107 |  | 0.645 |
| Threshold diff |  | - |  | Mood post AT |  | -0.108 |  | 0.641 |
| Threshold diff |  | - |  | Mood post CN |  | -0.061 |  | 0.792 |
| Threshold diff |  | - |  | Mood GB diff |  | 0.258 |  | 0.223 |
| Threshold diff |  | - |  | Mood AT diff |  | -0.215 |  | 0.313 |
| Threshold diff |  | - |  | Mood CN diff |  | -0.088 |  | 0.683 |
| Pain intensity diff |  | - |  | Mood pre GB |  | -0.361 |  | 0.084 |
| Pain intensity diff |  | - |  | Mood pre AT |  | -0.545 |  | 0.007 |
| Pain intensity diff |  | - |  | Mood pre CN |  | 0.027 |  | 0.901 |
| Pain intensity diff |  | - |  | Mood post GB |  | -0.684 | * | < .001 |
| Pain intensity diff |  | - |  | Mood post AT |  | -0.314 |  | 0.165 |
| Pain intensity diff |  | - |  | Mood post CN |  | -0.142 |  | 0.539 |
| Pain intensity diff |  | - |  | Mood GB diff |  | -0.459 |  | 0.025 |
| Pain intensity diff |  | - |  | Mood AT diff |  | -0.101 |  | 0.638 |
| Pain intensity diff |  | - |  | Mood CN diff |  | -0.128 |  | 0.550 |
| Pain unpleasantness diff |  | - |  | Mood pre GB |  | -0.211 |  | 0.320 |
| Pain unpleasantness diff |  | - |  | Mood pre AT |  | -0.461 |  | 0.025 |
| Pain unpleasantness diff |  | - |  | Mood pre CN |  | -0.179 |  | 0.401 |
| Pain unpleasantness diff |  | - |  | Mood post GB |  | -0.638 | * | 0.002 |
| Pain unpleasantness diff |  | - |  | Mood post AT |  | -0.413 |  | 0.064 |
| Pain unpleasantness diff |  | - |  | Mood post CN |  | -0.155 |  | 0.502 |
| Pain unpleasantness diff |  | - |  | Mood GB diff |  | -0.546 |  | 0.007 |
| Pain unpleasantness diff |  | - |  | Mood AT diff |  | -0.276 |  | 0.192 |
| Pain unpleasantness diff |  | - |  | Mood CN diff |  | -0.070 |  | 0.746 |
| * p < .004 | | | | | | | | |

Correlations between mood and placebo analgesia in VR in the analgesia-expectation group only. Spearman’s Rank correlations were performed to adjust for nonparametric data, and adjusted for multiple comparisons (new significance value = *P* < .004). ΔThreshold (Threshold diff), Δpain intensity (Pain intensity diff), and Δpain unpleasantness (Pain unpleasantness diff) were correlated with mood questions (GB: good-bad; AT: awake-tired; CN: calm-nervous) prior to beginning experiment (pre) and after completing the experiment (post), as well as with the difference in scores in mood between pre and post (diff). Better mood post experiment negatively correlated with pre to post intervention pain intensity [*r_s_*(12) = -0.68, *P* < .001] and pain unpleasantness ratings [*r_s_*(12) = -0.64, *P* < .01], meaning that higher ratings for good mood correlated to greater decreases from pre-intervention to post-intervention in pain intensity and unpleasantness. No other significant correlations were found (all comparisons *P* > .007, *r*(12) > -0.55).

Table 2 – Supplementary Materials.

| **Spearman Correlations: Mood and Placebo Analgesia in AR** | | | | | | | | |
| --- | --- | --- | --- | --- | --- | --- | --- | --- |
|  | |  | |  | | **Spearman's rho** | | **p** |
| Threshold diff |  | - |  | Mood pre GB |  | 0.051 |  | 0.812 |
| Threshold diff |  | - |  | Mood pre AT |  | 0.087 |  | 0.686 |
| Threshold diff |  | - |  | Mood pre CN |  | 0.281 |  | 0.184 |
| Threshold diff |  | - |  | Mood post GB |  | 0.213 |  | 0.354 |
| Threshold diff |  | - |  | Mood post AT |  | -0.013 |  | 0.955 |
| Threshold diff |  | - |  | Mood post CN |  | 0.122 |  | 0.598 |
| Threshold diff |  | - |  | Mood GB diff |  | 0.237 |  | 0.265 |
| Threshold diff |  | - |  | Mood AT diff |  | 0.053 |  | 0.807 |
| Threshold diff |  | - |  | Mood CN diff |  | 0.036 |  | 0.867 |
| Pain intensity diff |  | - |  | Mood pre GB |  | -0.144 |  | 0.499 |
| Pain intensity diff |  | - |  | Mood pre AT |  | -0.121 |  | 0.572 |
| Pain intensity diff |  | - |  | Mood pre CN |  | -0.540 |  | 0.007 |
| Pain intensity diff |  | - |  | Mood post GB |  | -0.364 |  | 0.106 |
| Pain intensity diff |  | - |  | Mood post AT |  | -0.269 |  | 0.238 |
| Pain intensity diff |  | - |  | Mood post CN |  | -0.290 |  | 0.202 |
| Pain intensity diff |  | - |  | Mood GB diff |  | -0.261 |  | 0.217 |
| Pain intensity diff |  | - |  | Mood AT diff |  | -0.205 |  | 0.334 |
| Pain intensity diff |  | - |  | Mood CN diff |  | 0.162 |  | 0.448 |
| Pain unpleasantness diff |  | - |  | Mood pre GB |  | -0.012 |  | 0.956 |
| Pain unpleasantness diff |  | - |  | Mood pre AT |  | 0.086 |  | 0.688 |
| Pain unpleasantness diff |  | - |  | Mood pre CN |  | -0.471 |  | 0.021 |
| Pain unpleasantness diff |  | - |  | Mood post GB |  | -0.383 |  | 0.087 |
| Pain unpleasantness diff |  | - |  | Mood post AT |  | -0.132 |  | 0.566 |
| Pain unpleasantness diff |  | - |  | Mood post CN |  | -0.312 |  | 0.169 |
| Pain unpleasantness diff |  | - |  | Mood GB diff |  | -0.331 |  | 0.114 |
| Pain unpleasantness diff |  | - |  | Mood AT diff |  | -0.114 |  | 0.595 |
| Pain unpleasantness diff |  | - |  | Mood CN diff |  | 0.137 |  | 0.520 |
| * p < .004 | | | | | | | | |

Correlations between mood and placebo analgesia in AR in the analgesia-expectation group only. Spearman’s Rank correlations were performed to adjust for nonparametric data, and adjusted for multiple comparisons (new significance value = *P* < .004). ΔThreshold (Threshold diff), Δpain intensity (Pain intensity diff), and Δpain unpleasantness (Pain unpleasantness diff) were correlated with mood questions (GB: good-bad; AT: awake-tired; CN: calm-nervous) prior to beginning experiment (pre) and after completing the experiment (post), as well as with the difference in scores in mood between pre and post (diff). No significant correlations that survived multiple comparisons were found (all comparisons *P* > .007, *r*(12) > -0.54).

| Table 3 – Supplementary Materials.  **Spearman Correlations: Mood and Placebo Analgesia in PR** | | | | | | | | |
| --- | --- | --- | --- | --- | --- | --- | --- | --- |
|  | |  | |  | | **Spearman's rho** | | **p** |
| Threshold diff |  | - |  | Mood pre GB |  | -0.362 |  | 0.082 |
| Threshold diff |  | - |  | Mood pre AT |  | -0.197 |  | 0.355 |
| Threshold diff |  | - |  | Mood pre CN |  | 0.185 |  | 0.387 |
| Threshold diff |  | - |  | Mood post GB |  | 0.129 |  | 0.547 |
| Threshold diff |  | - |  | Mood post AT |  | 0.432 |  | 0.035 |
| Threshold diff |  | - |  | Mood post CN |  | 0.303 |  | 0.182 |
| Threshold diff |  | - |  | Mood GB diff |  | 0.058 |  | 0.803 |
| Threshold diff |  | - |  | Mood AT diff |  | 0.283 |  | 0.214 |
| Threshold diff |  | - |  | Mood CN diff |  | 0.182 |  | 0.394 |
| Pain intensity diff |  | - |  | Mood pre GB |  | -0.199 |  | 0.352 |
| Pain intensity diff |  | - |  | Mood pre AT |  | -0.204 |  | 0.338 |
| Pain intensity diff |  | - |  | Mood pre CN |  | -0.065 |  | 0.762 |
| Pain intensity diff |  | - |  | Mood post GB |  | -0.219 |  | 0.302 |
| Pain intensity diff |  | - |  | Mood post AT |  | -0.148 |  | 0.489 |
| Pain intensity diff |  | - |  | Mood post CN |  | -0.196 |  | 0.393 |
| Pain intensity diff |  | - |  | Mood GB diff |  | -0.105 |  | 0.649 |
| Pain intensity diff |  | - |  | Mood AT diff |  | 0.138 |  | 0.550 |
| Pain intensity diff |  | - |  | Mood CN diff |  | -0.272 |  | 0.198 |
| Pain unpleasantness diff |  | - |  | Mood pre GB |  | -0.032 |  | 0.882 |
| Pain unpleasantness diff |  | - |  | Mood pre AT |  | 0.103 |  | 0.632 |
| Pain unpleasantness diff |  | - |  | Mood pre CN |  | -0.064 |  | 0.765 |
| Pain unpleasantness diff |  | - |  | Mood post GB |  | -0.041 |  | 0.850 |
| Pain unpleasantness diff |  | - |  | Mood post AT |  | -0.045 |  | 0.834 |
| Pain unpleasantness diff |  | - |  | Mood post CN |  | -0.256 |  | 0.262 |
| Pain unpleasantness diff |  | - |  | Mood GB diff |  | -0.126 |  | 0.585 |
| Pain unpleasantness diff |  | - |  | Mood AT diff |  | 0.014 |  | 0.953 |
| Pain unpleasantness diff |  | - |  | Mood CN diff |  | -0.387 |  | 0.063 |
| Threshold diff |  | - |  | Mood pre GB |  | -0.194 |  | 0.362 |
| Threshold diff |  | - |  | Mood pre AT |  | -0.025 |  | 0.908 |
| * p < .004 | | | | | | | | |

Correlations between mood and placebo analgesia in PR in the analgesia-expectation group only. Spearman’s Rank correlations were performed to adjust for nonparametric data, and adjusted for multiple comparisons (new significance value = *P* < .004). ΔThreshold (Threshold diff), Δpain intensity (Pain intensity diff), and Δpain unpleasantness (Pain unpleasantness diff) were correlated with mood questions (GB: good-bad; AT: awake-tired; CN: calm-nervous) prior to beginning experiment (pre) and after completing the experiment (post), as well as with the difference in scores in mood between pre and post (diff). No significant correlations that survived multiple comparisons were found (all comparisons *P* > .035, *r*(12) > -0.20).
